# Regulators of Interferon-Responsive Microglia Uncovered by Genome-wide CRISPRi Screening

**DOI:** 10.1101/2025.06.05.658176

**Authors:** Amanda McQuade, Vincent Cele Castillo, Venus Hagan, Weiwei Liang, Thomas Ta, Reet Mishra, Olivia Teter, Noam Teyssier, Kun Leng, Martin Kampmann

**Affiliations:** Institute for Neurodegenerative Diseases, University of California, San Francisco, San Francisco, CA, USA; UC Berkeley-UCSF Graduate Program in Bioengineering, University of California, San Francisco, San Francisco, CA, USA; Biomedical Informatics Graduate Program, University of California, San Francisco, San Francisco, CA, USA; Medical Scientist Training Program, University of California, San Francisco, San Francisco, CA, USA; Biomedical Sciences Graduate Program, University of California, San Francisco, San Francisco, CA, USA; Department of Biochemistry and Biophysics, University of California, San Francisco, San Francisco, CA, USA

## Abstract

Microglia dynamically support brain homeostasis through the induction of specialized activation programs or states. One such program is the Interferon-Responsive Microglia state (IRM), which has been identified in developmental windows, aging, and disease. While the functional importance of this state is becoming increasingly clear, our understanding of the regulatory networks that govern IRM induction remain incomplete. To systematically identify genetic regulators of the IRM state, we conducted a genome-wide CRISPR interference (CRISPRi) screen in human iPSC-derived microglia (iPS-Microglia) using IFIT1 as a representative IRM marker. We identified 772 genes that modulate IRM, including canonical type I interferon signaling genes (*IFNAR2, TYK2, STAT1/2, USP18*) and novel regulators. We uncovered a non-canonical role for the CCR4-NOT complex subunit CNOT10 in IRM activation, independent of its traditional function. This work provides a comprehensive resource for dissecting IRM biology and highlights both established and novel targets for modulating microglial interferon signaling in health and disease.

## Introduction

Microglia are dynamic immune cells that continuously monitor the brain parenchyma to support brain health. In response to insult or injury, microglia rapidly trigger transcriptional activation programs to restore homeostasis. Although microglial immune responses have been traditionally thought of as merely a response to disease pathology, the last decade of research has centered microglia as a disease-modifying cell type across many developmental and neurodegenerative conditions.

One microglial activation state that shows clear functional impact on brain health is the Interferon-Responsive Microglia (IRM) state, defined by high expression of interferon-stimulated genes. Type I Interferons (IFN), including IFNα, IFNβ, IFNε, IFNκ, and IFNϖ, bind to type 1 interferon receptors, IFNAR1/2, and induce a JAK/STAT signaling cascade, leading to nuclear translocation of the transcription factor complex ISGF3 containing STAT1, STAT2, and IRF9, and transcription of interferon-stimulated genes such as IFIT1. These genes reinforce interferon signaling through autocrine and paracrine loops. While interferon responses were originally characterized as a mechanism of host defense against viral particles, microglia have been found to strongly induce ISGs even in the absence of viral infection^1–4^. The source of these IFN responses may differ in developmental, aging, or disease paradigms. IFN responses can be induced by detection of double-stranded RNA elements (such as microRNAs) or DNA even in sterile conditions. These interferon-relevant stimuli may build up in the brain during aging as mitochondrial DNA leakage becomes more prevalent or removal of apoptotic cellular debris becomes less efficient. Microglia may also detect interferon in a paracrine fashion. For example, the choroid plexus has been shown to increase secretion of IFNβ into the parenchyma particularly in aging^1^, which would impact microglial activation.

In developmental windows, IRM play critical roles in synaptic refinement^5^. Interestingly, heightened interferon responses have been identified in Autism Spectrum Disorder (ASD) patients^6^ and in models of ASD *de novo* mutations^7^. However, in normal development, IRM activation is tightly temporally regulated, and this activation state does not persist into healthy adulthood. In the aging brain, IRM expand again, with over half of microglial aging genes being related to Type I interferon responses^8,9^. In recently established brain aging clocks, nearly all brain cell types show a strong interferon response signature in the aged brain, which may be linked to the cognitive decline associated with aging^10^. IRM have also been identified to be further enriched in neurodegenerative diseases such as Alzheimer’s disease^2,4,11^.

Despite the identification of IRM across aging and disease datasets, it is still not fully understood if IRM represent a beneficial activation program, working to combat aging and disease phenotypes, or if IRM contribute to disease progression. The impact of IRM is likely to be disease-, pathology-, and temporally-specific. Current evidence suggests that increased interferon signaling in the brain enhances microglial phagocytosis, potentially inducing aberrant synaptic pruning, which leads to dementia-like symptoms^2,4,12^. Blocking interferon responses with transgenic mice lacking the type I interferon receptor (IFNAR1), anti-IFNAR antibodies, or cGAS-STING inhibitors that reduce IRM, has shown efficacy in slowing cognitive decline in normal aging contexts as well as in Alzheimer’s disease models, driven by either amyloid or tau, and in Parkinson’s disease^4,13–17^. However, IFNβ and IRM are not always detrimental. For example, IFNβ has been used to treat patients with multiple sclerosis^18,19^. In Experimental Autoimmune Encephalomyelitis (EAE) models of Multiple Sclerosis, activation of microglial interferon responses, specifically through ganciclovir treatment, proved therapeutic. Similarly, IFNβ stimulation promoted beneficial microglia functions, including phagocytosis of α-synuclein and autophagic flux, in Parkinson’s disease models^20^.

Taken together, observations of IRM across development, aging, and disease point to complex regulatory processes that underlie induction of IRM. This motivated us to systematically evaluate genetic regulators of the IRM state with a genome-wide CRISPR-interference screen. Although the direction of IRM regulation for therapeutic purposes may be context-specific, this dataset provides a foundation to better understand the functions of IRM or develop therapeutic targets to perturb microglial activation. We recovered a large proportion of canonical interferon signaling regulators and ISGs that regulate microglial access to the IRM state. Additionally, we identified novel regulators of IRM, including a non-canonical role of the CCR4-NOT deadenylase subunit CNOT10.

## Results

### Markers of the interferon response in iPSC-derived microglia

To enable a genome-wide screen for modifiers of the type I interferon response in induced pluripotent stem cell-derived microglia (iPS-Microglia), we first identified IFNβ-responsive transcripts via bulk RNA-sequencing in this microglia model. We treated iPSC-microglia with low (10 pM) or high (100 pM) concentrations of IFNβ (Figure 1A,B, Supplemental Figure 1, Supplemental Table 1). We found significant correlation of differentially expressed genes (DEGs) (R^2^=0.93) in these two treatment concentrations compared to untreated iPS-Microglia, although globally, 100 pM IFNβ treatment drove larger effect sizes and resulted in a greater number of DEGs than 10 pM IFNβ (Supplemental Figure 1).

**Figure 1.**
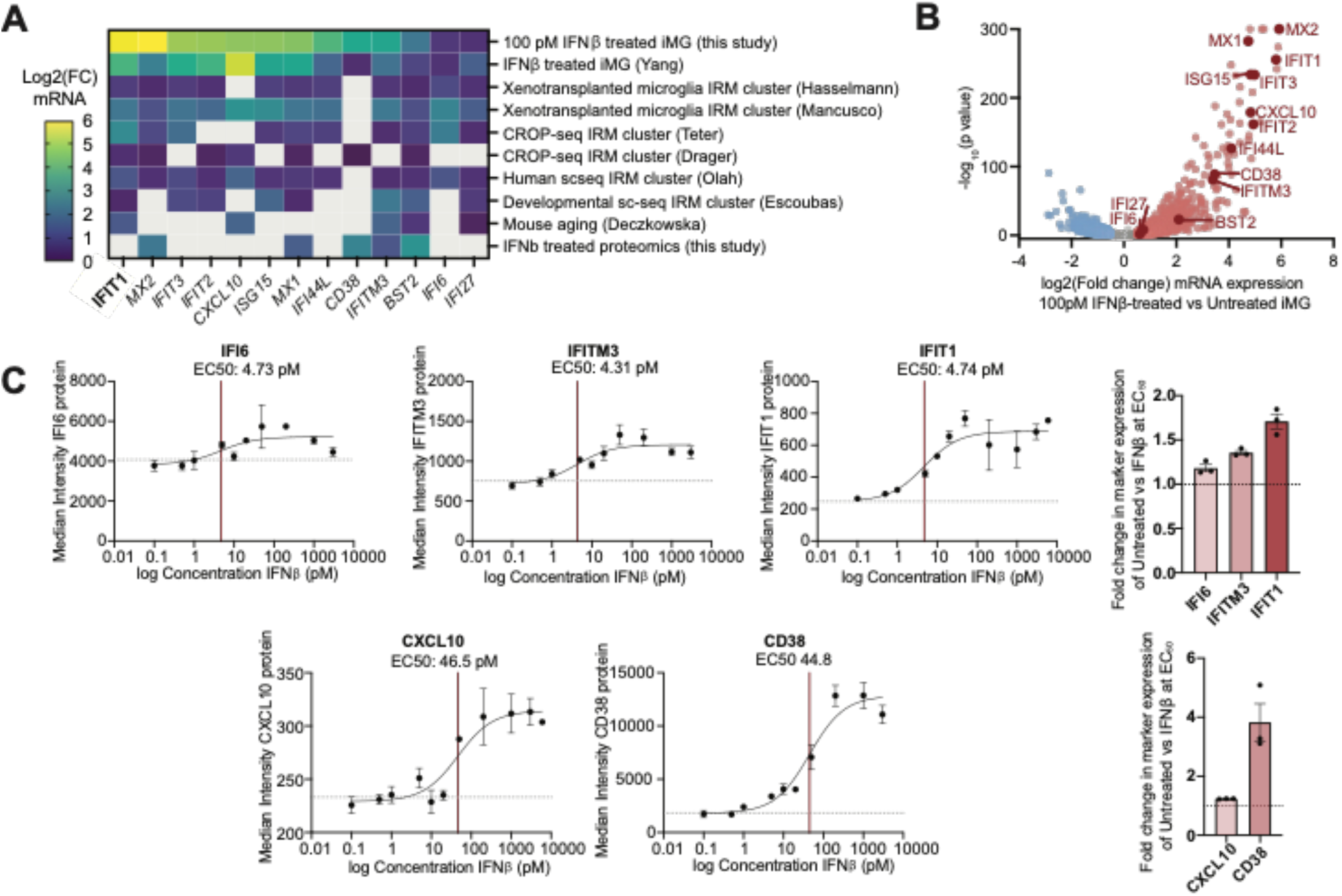
Identification of robust markers for IRM. **(A)** Comparison of mRNA expression across models of IRM. Log2(FC) mRNA expression for 100 pM IFNβ vs untreated iPS-Microglia (bulk RNA-seq, this study), 100 U/mL IFNβ vs untreated iPS-Microglia (bulk RNA-seq, Yang), IRM cluster versus all other clusters (Single-cell seq, Hasselmann), IRM cluster vs all other clusters (Single-cell seq, Mancuso), IRM cluster versus all other clusters (CROP-seq, Teter), IRM cluster versus all other clusters (CROP-seq, Drager), IRM cluster versus all other clusters (Single-cell seq, Olah), IRM cluster versus all other clusters (Single-cell, Escoubas), 3 month microglia verus 22 month old microglia (bulk RNA-seq, Deczkowska). Grey fill represents that the given gene is not significantly altered in the dataset. **(B)** Volcano plot showing differentially expressed genes in 100 pM IFNβ treated iPS-Microglia versus untreated. Conserved ISGs are denoted in dark red. **(C)** Dose response curves for IFI6, IFITM3, IFIT1, CXCL10, and CD38. Median fluorescence intensity (MFI) of IFIT1 was measured after 24 hours of treatment with IFNβ. Dotted horizontal line represents MFI in untreated iPS-Microglia. Red vertical line represents the IFN concentration that yields 50% MFI response. Fold change of MFI 5 pM (top) or 50 pM (bottom) shown on right. Graphs represent mean +/-standard error. n=3 independent wells, >10,000 cells per well (individual well datapoints shown on right)

To identify the most robust markers of interferon-responsive microglia (IRM), we cross-referenced our interferon-responsive transcripts against several published datasets, focusing on those that used human postmortem tissue or human iPSC-derived models^5,7,9,21–25^ (Figure 1A). Five IRM marker proteins from this subset were selected as candidate markers for a genome-wide screen. For each protein, response to IFNβ was measured by flow cytometry over a broad range of IFNβ concentrations (Figure 1C). Interestingly, three candidates (IFI6, IFITM3, IFITM1) had an EC^50^ of ∼5 pM IFNβ, whereas two candidates (CXCL10, CD38) had an EC^50^ of ∼50 pM. This mirrors a previous single-cell sequencing analysis we performed on iPS-Microglia, which revealed two distinct IRM clusters, only one of which showed elevated CXCL10 and CD38^25^. Because the markers that responded at ∼5 pM IFNβ were more conserved across datasets we wanted to maximize discovery of this class of interferon-response gene. Interferon-Induced Protein with Tetratricopeptide Repeats 1 (IFIT1) showed the best dynamic response range and was chosen for our genome-wide screen. IFIT1 has been identified as a specific marker of the type I IFN-response pathway, lending further specificity to our screening results^26^.

To determine regulators of IRM, we optimized a protocol (see Methods) to sort iPS-Microglia into high versus low interferon responders (top or bottom 30% IFIT1 expression) (Figure 2A, Supplemental Table 2). Using an empirical discovery rate of 5% based on non-targeting control sgRNAs (NTC), we identified 772 genes for which knockdown regulated the interferon response (Figure 2B). For 500 genes, targeted CRISPR-interference decreased levels of IFIT1. For 272 genes, CRISPR-interference promoted levels of IFIT1. Top hits for which knockdown (KD) negatively regulated interferon response include canonical interferon signal transduction pathway effectors (*TYK2, IFNAR2, JAK1, STAT1, STAT2, IRF9*) (Figure 2B,C). Canonical negative regulators of interferon signal transduction were identified as permitting higher interferon response after knockdown (*ISG15, USP18)*. Interestingly, knockdown of homeostatic signaling receptors (*TGFBR1, TGFBR2)* also promoted IFIT1 levels after IFNβ stimulation, suggesting that these proteins normally act to attenuate IFNβ signaling.

**Figure 2.**
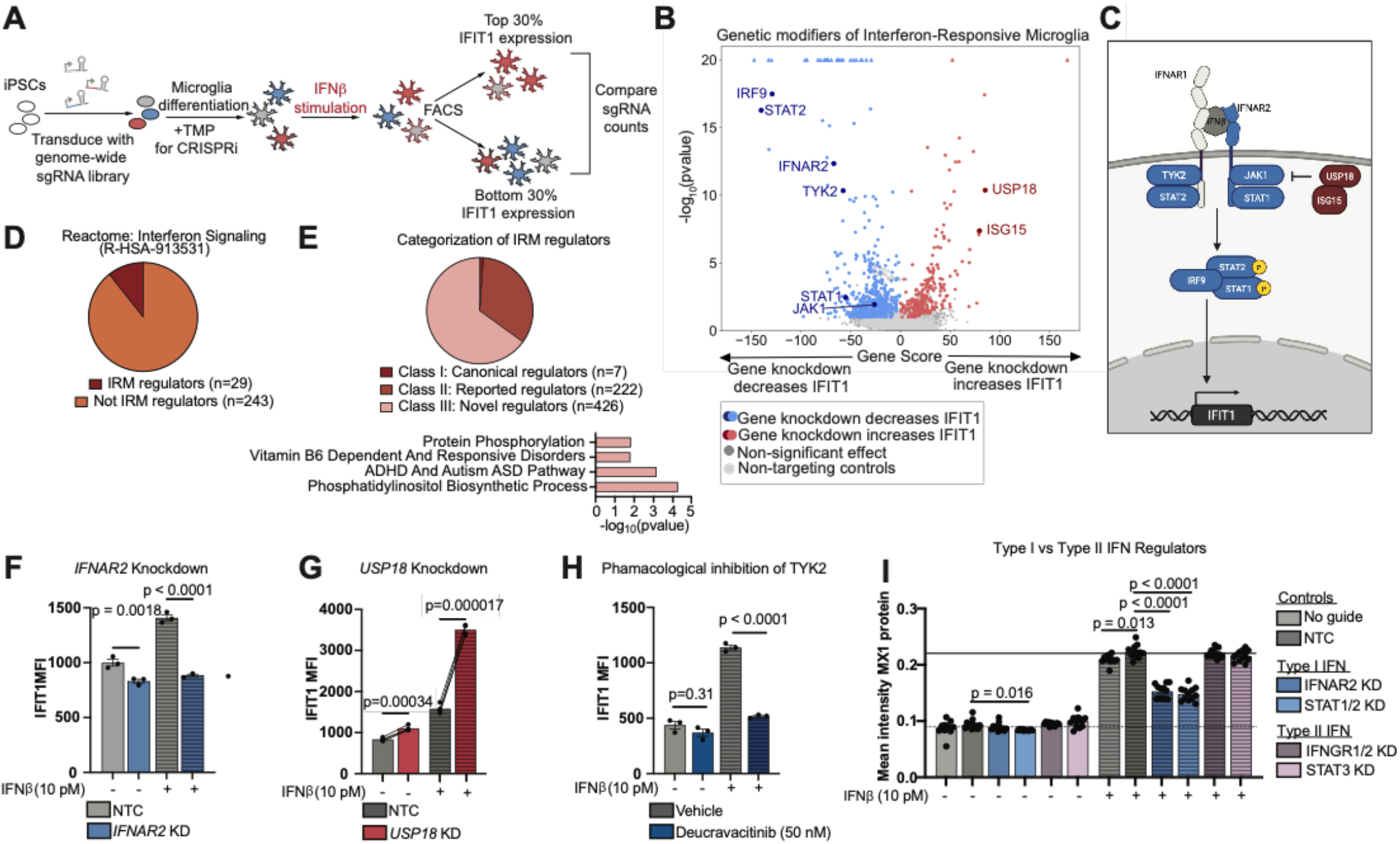
Interferon-responsive microglia are responsive to canonical interferon regulators. **(A)** Schematic of genome-wide screen for IFIT1 expression. **(B)** Volcano plot of results from genome-wide screen. Gene score (normalized log2FC of guide counts in IFIT1 high versus IFIT1 low populations) is plotted versus the negative log of the p value. Each dot represents Gene score for a single gene averaged across five individual sgRNAs. Positive hits (gene knockdown drives IFIT1) are plotted in pink, negative hits (gene knockdown inhibits IFIT1) are plotted in blue. Non-targeting controls are in light grey and non-hit genes are in grey. Canonical interferon pathway genes are labelled. **(C)** Schematic of type I interferon signaling pathway. Positive hits are shown in red and negative hits are shown blue, non-hits are shown in grey. Blue and grey colored genes make up the Google Gemini canonical interferon response genes. **(D)** Overlap of screen hits with the Reactome “Interferon Signaling” classification. **(E)** Google Gemini classification of screen hits (see Methods) as canonical interferon regulators (darkest red, these genes are also shown in Figure 2C in blue and grey), genes reported to be associated with interferon regulation (red) or novel interferon regulators (light red). Gene ontology biological processes and wiki pathways analysis for the novel descriptors is shown below. pvalue of enrichment calculated by ENRICHR. **(F)** Mean fluorescence intensity (MFI) of IFIT1 protein levels in IFNAR2 knockdown (blue) versus non-targeting control (NTC, grey). 10 pM IFNβ was added for 24 hrs before flow cytometry analysis. (n=3 technical replicates, >10,000 cells per replicate. Bars represent mean +/-standard error, T test) **(G)** Mean fluorescence intensity (MFI) +/-standard error of IFIT1 protein levels in USP18 knockdown (red) versus non-targeting control (NTC, grey) samples plated as in-well controls. 10 pM IFNβ was added for 24 hrs before flow cytometry analysis. (n=3 technical replicates, >10,000 cells per replicate. Bars represent mean +/-standard error, paired T test). **(H)** Mean fluorescence intensity (MFI) of IFIT1 protein levels after Deucravacitinib (blue) versus vehicle (grey). After 30 min pre-treatment with Deucravacitinib or vehicle, 10 pM IFNβ was added for 24 hrs before flow cytometry analysis. (n=3 technical replicates, >10,000 cells per replicate. Bars represent mean +/-standard error, T test compared to non-targeting control). **(I)** Mean fluorescence intensity (MFI) of MX1 protein levels normalized to DAPI nuclear counts after knockdown versus non-targeting control. 10 pM IFNβ was added for 24 hrs before fixation. (n= 4 independent wells averaging from 4 images per well, Bars represent mean +/-standard error, ANOVA with Tukey’s multiple comparison test).

Comparing our gene set of IRM regulators to the Reactome term *Interferon Signaling* revealed limited overlap (29/272 genes, Figure 2D). However, we did find enrichment of predicted STAT1 and STAT2 regulons (TRRUST v2; FDR STAT1: 0.0001, STAT2: 0.0357) for the negative hits and enrichment of SP1 and IRF2 in the positive hits (TRRUST v2; FDR SP1: 0.0084, IRF2: 0.0212). As interferon regulation in peripheral systems has been under focused research for decades, we mined existing research studies using the Google Gemini large language model. Google Gemini was used to classify our screen hits into “canonical interferon regulators”, which include those directly downstream of IFNβ (Figure 2C, labelled in blue and grey), genes with a “reported association with interferon”, which include genes found in IFN- and viral-response gene ontology terms, as well as those for which one or more publications mention the gene in the context of IFN, and “novel interferon regulators”, for which Google Gemini did not find any known associations with IFN response (Figure 2E, Supplemental Table 2, Supplemental Table 3,4). For the novel interferon regulators revealed by our screen, the following gene ontology and wikipathway terms were enriched: *GO:0006468 Protein Phosphorylation, WP4228 Vitamin B6 Dependent and Responsive Disorders, WP5420 ADHD and Autism ASD Pathway*, and *GO:0006661 Phosphatidylinositol Biosynthetic Process* (Figure 2E).

### Canonical regulation of interferon is relevant in iPS-Microglia models

To validate our screening results, we generated independent knockdown lines of canonical interferon pathway hits (Supplemental Figure 2). Knockdown of type I interferon receptor, *IFNAR2*, decreased IFIT1 levels at baseline suggesting this model of iPS-Microglia has some level of tonic IFNβ production. This corroborates previous findings, highlighting the existence of IRM at baseline by single-cell RNA sequencing analyses^7,25^. As expected *IFNAR2* KD microglia show no increase in IFIT1 expression after treatment with IFNβ (Figure 2F).

Ubiquitin Specific Peptidase 18 (USP18) competitively inhibits JAK1 binding to IFNAR2 and blocks the necessary dimerization of IFNAR1 and IFNAR2. Knockdown of this negative interferon regulator increased IFIT1 levels both at baseline and after interferon stimulation (Figure 2G). Knockdown of USP18 increased iPS-Microglia response to IFNβ by over twofold, suggesting that USP18 may be a useful node to drive IFN responses therapeutically.

Tyrosine Kinase 2 (TYK2) associates with IFNAR1/2 to phosphorylate the Signal Transducer and Activator of Transcription 1 and 2 (STAT1/2) transcription factors that are necessary for transcription of interferon-responsive genes, including IFIT1. As expected, in our iPS-Microglia system, IFNβ-induced expression of IFIT1 is blocked by the TYK2 inhibitor, Deucravacitinib, though no difference in IFIT1 levels was detected in the absence of IFNβ stimulation (Figure 2H).

Both Type I and Type II interferon have been implicated in the progression of Alzheimer’s disease, with plaque-associated microglia showing high levels of IFIT1 and MX1^2,13,27^. Using CRISPRi knockdown of canonical Type I and Type II interferon signal transduction genes, we showed that MX1 is also responsive to IFNβ stimulation and that this could be specifically blocked with knockdown of Type I IFN pathway genes (*IFNAR2, STAT1/2*) but not Type II IFN (IFNGR1/2, STAT3) (Figure 2I).

### CCR4-NOT Transcription Complex Subunit 10 (CNOT10) knockdown inhibits IRM

Two of our most significant screen hits where knockdown decreases IRM are CCR4-NOT Transcription Complex Subunit 10 and 11 (CNOT10, CNOT11). CNOT10 and CNOT11 are both subunits of the CCR4-NOT transcription complex that is responsible for deadenylating mRNA (Figure 3A). Interestingly, none of the enzymatic deadenylase subunits (*CNOT6, CNOT6L, CNOT7, CNOT8)* or the scaffold of the complex (*CNOT1)* were discovered as IRM regulators by our IRM screen. Indeed, in independent knockdown lines, only *CNOT10* KD decreased IFIT1 expression (Figure 3B, C, Supplemental Figure 2), suggesting that the canonical deadenylation functions of this complex do not explain the difference in interferon response seen in *CNOT10* KD microglia. We confirmed that *CNOT10* knockdown inhibited IRM using multiple markers of the state and a wide range of concentrations (Figure 3C,D). Our data suggests that *CNOT10* KD microglia have a maximal response to interferon that is lower than their WT counterparts and a left-shifted EC^50^ of IFIT1 response.

**Figure 3.**
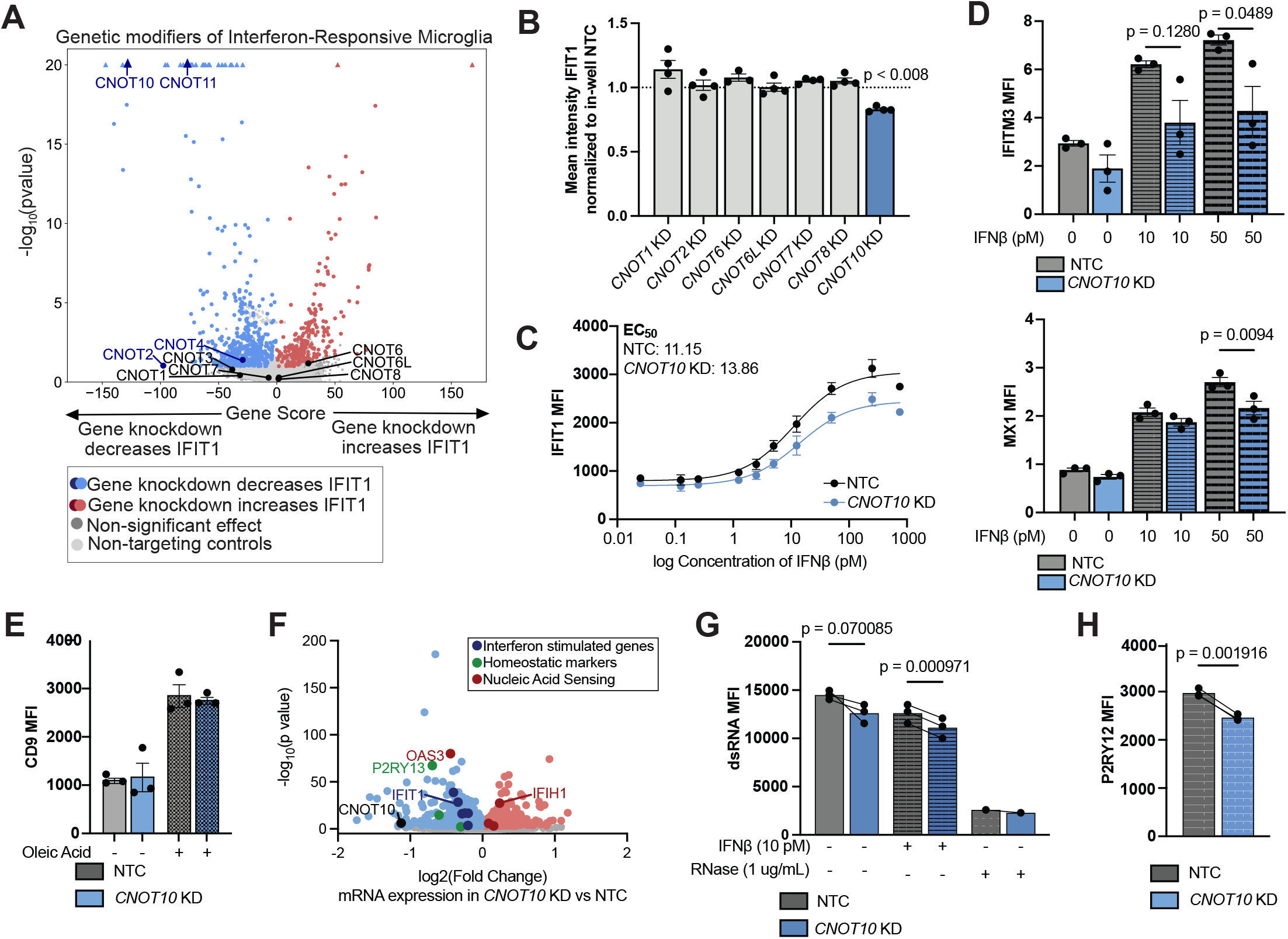
Knockdown of CNOT10 inhibits IRM. **(A)** Volcano plot of results from genome-wide screen. Gene score (normalized log2FC of guide counts in IFIT1 high versus IFIT1 low populations) is plotted versus the negative log of the p value. Each dot represents Gene score for a single gene averaged across five individual sgRNAs. Positive hits (gene knockdown drives IFIT1) are plotted in pink, negative hits (gene knockdown inhibits IFIT1) are plotted in blue. Non-targeting controls are in light grey and non-hit genes are in grey. All members of the CCR4-NOT complex are labelled (significant genes in blue, non-significant genes in black). **(B)** Mean fluorescence intensity (MFI) of IFIT1 protein levels in CCR4-NOT family normalized to non-targeting in-well controls (n=3 technical replicates, >10,000 cells per replicate. Bars represent mean +/-standard error, T test). **(C)** Dose response curves for interferon response in CNOT10 knockdown (blue) vs non-targeting control (NTC, grey). Median fluorescence intensity (MFI) of IFIT1 was measured after 24 hours of treatment with IFNβ. Data represent mean +/-standard error. n=3 independent wells >10,000 cells per well. EC50 calculated as nonlinear fit [agonist] vs response in Prism. **(D)** Mean fluorescence intensity (MFI) of IFITM3 (top) and MX1 (bottom) protein levels in CNOT10 knockdown (blue) versus non-targeting control (NTC, grey). IFNβ was added at the indicated concentration for 24 hrs before flow cytometry analysis. (n=3 technical replicates, >10,000 cells per replicate. Bars represent mean +/-standard error, ANOVA, Tukey’s multiple comparison test). **(E)** Mean fluorescence intensity (MFI) CD9 protein levels in CNOT10 knockdown (blue) versus non-targeting control (NTC, grey). 250 nM Oleic acid was added 24 hrs before flow cytometry analysis. (n=3 technical replicates, >10,000 cells per replicate. Bars represent mean +/-standard error, no significance, ANOVA, Tukey’s multiple comparison test). **(F)** Volcano plot of differentially expressed genes in CNOT10 KD versus non-targeting control (NTC). Genes that increase or decrease in expression in CNOT10 KD are colored red or blue (FDR < 0.05). Genes related to interferon, microglia homeostatic markers, or nucleic acid sensing are highlighted. **(G)** Double-stranded RNA (dsRNA) was measured by flow cytometry in iPS-Microglia with *CNOT10* knockdown or in-well, non-targeting control sgRNA. For the indicated bars, 10 pM IFNb was added for 24 hrs or 1 ug/mL RNase A for 30 min after fixation. Bars represent mean fluorescence intensity +/-standard error, Paired t-test. For NTC and *CNOT10* KD comparison, n=3 independent wells, >10,000 cells per well. For RNAse control, n=1 well, >10,000 cells per well. **(H)** Mean fluorescence intensity (MFI) P2RY12 protein levels in CNOT10 knockdown (blue) versus non-targeting control (NTC, grey). (n=3 technical replicates, >10,000 cells per replicate. Bars represent mean +/-standard error, paired T test).

Because CNOT10 is part of a transcription regulation complex and could be acting via broad mRNA destabilization, we tested the specificity of the inhibition response. Upon stimulation with oleic acid, both non-targeted and *CNOT10* knockdown iPS-Microglia activated CD9 expression to the same extent (Figure 3E). To more closely examine the impact of *CNOT10* knockdown on microglial activation and interferon-response, we performed single-cell RNA-sequencing on iPS-Microglia expressing a *CNOT10* knockdown sgRNA or non-targeting sgRNA after 24 hrs of IFNβ stimulation (Figure 3F, Supplemental Table 5). In support of our hypothesis that the inhibition of IRM is not due to generalized mRNA deadenylation, we found approximately equal numbers of differentially expressed genes in the positive and negative direction (403 increased expression, 506 decreased expression in *CNOT10* KD).

Corroborating our quantification at the protein level, mRNA levels of multiple interferon-stimulated genes were decreased after *CNOT10* knockdown, specifically *IFIT1, IFIT3, IFI6, IFI30, IFIT5*, and *IFITM1* (Figure 3F). We also lose expression of *S100B* in *CNOT10* knockdown microglia, a secreted factor that has recently been shown to be critical for binding of IFNβ to IFNAR1/2^28^. Surprisingly, transcript levels of nucleic acid sensors Cyclic GMP-AMP Synthase (*CGAS)* and Interferon-Induced with Helicase C Domain 1 (*IFIH1)*, encoding MDA5 were increased in *CNOT10* KD microglia. However, the upstream dsRNA-binding protein 2’, 5’ oligoadenylate (*OAS3)* was decreased in expression in *CNOT10* KD microglia. Additionally, we did not find evidence of increased dsRNA in *CNOT10* knockdown microglia (Figure 3G).

Further corroborating our protein level analysis of *CNOT10* knockdown, our RNA-sequencing data shows no difference in mRNA expression of *CD9*. However, despite *CNOT10* KD microglia not seeming robustly activated, our RNA-sequencing data revealed lower levels of microglia homeostatic markers, *CX3CR1, P2RY12* and *P2RY13* (Figure 3F). We additionally confirmed P2RY12 to be decreased at the protein level (Figure 3H), which may suggest that CNOT10 is altering microglial surveillance.

Importantly, expression of *CNOT10* is decreased at the mRNA level in our *CNOT10* knockdown lines, and expression of all other CCR4-NOT complex subunits remain unchanged (Supplemental Table 5). Since the CCR4-NOT complex can function without CNOT10^29,30^, it is likely that our knockdown lines remain competent in deadenylation. However, *CNOT10* knockdown microglia showed changes to gene expression that could impact global translation. Genes encoding subunits of both the small and large ribosomal proteins were enriched among mRNAs decreasing in *CNOT10* knockdown microglia. On the other hand, many translation initiation factors were increased in expression in the *CNOT10* knockdown microglia. TATA-box binding protein associated factor, RNA polymerase I subunit B (*TAF1B*) was also increased in expression in *CNOT10* knockdown cells. Together, these data suggest a disruption to translation, though it remains unclear how this would impact specific microglial activation states (IRM, Homeostatic) and not others (DAM) as our data suggests.

## Discussion

In this study, we present a genome-wide CRISPR-interference screen for expression of the Type I interferon marker IFIT1. An initial investigation of the concentration-dependent responses of interferon-stimulated genes (ISGs) at the transcriptomic level revealed high correlation between ISG induction at 10 pM vs 100 pM, suggesting that there are not two distinct response profiles at the two concentrations. However, at the protein level, a subset of ISGs including CXCL10 and CD38 showed a median response at 50 pM, whereas the majority of ISGs peaked at a lower IFNβ concentration (∼5 pM). In future work, it may prove useful to understand the functional differences between low IFN-responsive profiles and high IFN-responsive profiles to determine if there are two classes of interferon-responsive microglia or if higher concentrations merely exacerbate the same functions.

The data generated from our genome-wide screen supports the hypothesis that IRM are regulated via canonical interferon signal transduction. This finding suggests tools developed to combat peripheral interferon- and viral-responses can be translated for use in the central nervous system. For example, we highlight that knockdown of *STAT1, STAT2, IFNAR2*, and *USP18* modulate IFIT1 expression in microglia in the expected direction. Additionally, we show that treatment with Deucravacitinib, the FDA-approved inhibitor of TYK2, blocks interferon responses in iPS-Microglia. Moving forward, our full catalog of IRM regulators can be used to nominate other existing, FDA-approved therapeutics that could perturb microglial interferon responses.

In addition to the identification of canonical interferon-signaling genes, our screen has revealed a large number of novel regulators of IRM as well. Gene ontology terms of ADHD and Autism spectrum disorder were enriched in this population. Specifically, genes for which decreased expression or loss of function is correlated with ASD were hits in our screen; their knockdown inhibited interferon-responsive microglia (*GABBR1, ASMT, KCNA4, OPRM1, GLS, GATM, GRIN3A*). Future research should investigate the hypothesis that these loci impact ASD risk through perturbation of IRM.

Similarly, there was significant overlap between our screen hits and genes in AD risk loci. From 1077 genes in AD risk loci, 128 were hits in our genome-wide screen. For 101 genes, targeting with CRISPRi drives interferon-response and for 27, targeting with CRISPRi inhibits interferon response. However, it remains unclear what this means for Alzheimer’s pathology, since many of the AD risk loci have not been mapped to a particular causal gene and even for those with causal genes proposed, the functional impact of the SNP is not well understood. However, of particular interest, we found that knockdown of ADAM10 decreases IRM, while knockdown of APOE or INPP5D drives IRM. Previous studies have shown that APOE ε4 variant carriers have higher levels of IFNβ in the serum and plasma^11^. Likewise, APOE ε4 itself has been identified as a ligand for Leukocyte Immunoglobulin-Like Receptor B3 (LilrB3) which triggers a downstream interferon response whereas APOE ε2 does not produce this response^31^. Similarly, INPP5D has also been recently shown to negatively regulate interferon through binding of IRF3 in models of viral infection. Whether this mechanism underlies the impact of INPP5D on neurodegenerative disease remains unclear.

Two of our top novel regulators of IRM are CCR4-NOT Transcription Complex subunits 10 and 11 (CNOT10, CNOT11). CNOT10 and CNOT11 were identified in 2013 as novel subunits of the eukaryotic CCR4-NOT complex. These subunits evolved more recently than the rest of the CCR4-NOT complex and have no orthologues in yeast^32^. While CNOT10 and CNOT11 do contribute to canonical functions of CCR4-NOT deadenylation, particularly helping with ribosomal binding^33^, CNOT10 and CNOT11 have also been shown to interact with Gametogenetin Binding Protein 2 (GGNBP2)^32,34^. This interaction is proposed to influence nucleic acid recognition pathways by re-localizing dsRNA that has not been edited by Adenosine Deaminase RNA-Specific (ADAR) away from the cytoplasm, resulting in decreased interferon response^34^. Heraud-Farlow and colleagues show that knockdown of *CNOT10* or *CNOT11* in myeloid cells limits interferon response, which directly matches the effect we observed. However, Heraud-Farlow et al. only find this interaction to occur in ADAR editing mutants, while our iPS-Microglia are ADAR competent. Additionally, we do not detect substantial changes in dsRNA levels in our *CNOT10* knockdown microglia.

In opposition, Gordon et al. found *CNOT10* knockdown amplifies interferon responses, leading to more potent antiviral activity. This study was performed with HeLa cells and primary CD4+ T cells, which may explain the difference in findings^35^. It is possible that the effects we and Heraud-Farlow have found are specific to the myeloid lineage, though further research is required to address this hypothesis.

In summary, our results define a catalog of canonical and noncanonical regulators of the interferon-responsive microglial activation state and point to nodes within the interferon-response pathway that could be harnessed to tune microglia function across diverse developmental and disease contexts.

## Methods

### iPSC maintenance

Human iPSCs (WTC11 background with doxycycline-inducible transcription factors described below^25^) were cultured in StemFlex (Gibco, A33493-01) on Matrigel (Corning, 356231) coated plates. Media was changed every day and cells were passaged once per week using ReLeSR (100-0483). Human iPSC studies at the University of California, San Francisco were approved by the Human Gamete, Embryo and Stem Cell Research Committee.

### Differentiation of iPS-Microglia

iPS-Microglia were differentiated as previously described^25^ with a modified media composition optimized for increased expression of P2RY12, TREM2, and CD33. Induced pluripotent stem cells containing six doxycycline-inducible transcription factors: CEPBA, CEBPB, IRF5, IRF8, MAFB, PU.1, were passaged as a single cell suspension using accutase (Thermo Fisher Scientific, A11105-01) for 7 minutes at 37 C. Cells are plated to dual coated Matrigel (Corning, 356231) and PDL plates (Corning, 356470) and cultured for 48 hours in Essential 8 (Gibco, A1517001) stem cell maintenance media supplemented with 2 ug/mL doxycycline (Clontech, 631311) and 10 nM Y-27632 (Tocris 1254). Two days post passage, cells are moved to microglia differentiation media: BrainPhys (StemCell Technologies, 05790), 0.5x N2 Supplement (Gibco, 17502048), 0.5x B27 (Gibco, 17504044), 10 ng/mL NT-3 (Peprotech, 450-03), 10 ng/mL BDNF (Peprotech, 450-02), 1 ug/mL mouse laminin (Thermo Fisher Scientific, 23017-015) supplemented with 2 ug/mL doxycycline, 100 ng/mL IL-34 (Peprotech 200-34), and 10 ng/mL GM-CSF (Peprotech 300-03). On day 4, day 8, and day 12, media was replaced with microglia differentiation media supplemented with 2 ug/mL doxycycline, 100 ng/mL IL-34, 10 ng/mL GM-CSF, 50 ng/mL M-CSF (Peprotech 300-25), and 50 ng/mL TGFB (Peprotech100-21C). For experiments with CRISPRi, cells were supplemented with 50 nM TMP (MP Biomedical, 195527) every other day to maintain strong knockdown.

### Flow cytometry

After 13 days of differentiation, iPS-Microglia are lifted with TrypLE (Gibco, 12604021) for 10 min at 37 C. Enzymatic activity is quenched with equal volumes of Advanced DMEM F12 (Gibco, 12634028) and cells are pelleted at 300 xG for 5 min. For intracellular staining (IFIT1, IFITM3, IFI6, CXCL10, MX1, dsRNA) cells are fixed with eBioscience Intracellular Fixation and Permeabilization Buffer Set (Invitrogen, 888-8824-00) for 20 min at room temperature. After washing with permeabilization buffer, antibodies are added for 30 min at 4C in permeabilization buffer (IFIT1, Cell Signaling, 20329S, 1:100; IFITM3, Proteintech, 11714-1-AP, 1:100; IFI6, Invitrogen, PA5-76860, 1:100; CXCL10, Biolegend, 519504, 1:50; dsRNA, Absolute Antibody, Ab00458-23.0, 1:250). For dsRNA, cells are washed and secondary antibody Goat anti-Rabbit 488 (Invitrogen, A-11008, 1:500) is added for 30 min. For extracellular markers (CD38, CD9, P2RY12), cells were not fixed and incubated with antibodies for 30 min at 4C (FC block, Biolegned, 422302, 1:200; CD38, R&D Systems, FAB2404P, 1:200; CD9, Biolegend, 312104, 1:200; P2RY12, Biolegend, 392108, 1:50).

Samples were analyzed on a BD LSR Fortessa X14 using BD FACSDiva software. Mean fluorescence intensity was calculated using FlowJo analysis software after gating for live, single cells. Data was collected from 3 independent differentiations with 3 replicate wells per experiment (n > 10,000 cells).

For stimulated conditions, IFNβ (GIbco, AF-300-02B) was added at 10 pM unless otherwise indicated 24 hrs before lifting. For analysis of secreted proteins (CXCL10) GolgiPlug (BD Biosciences, 555029, 1:1000) was added 6 hours prior to lifting. For analysis of dsRNA, cells were treated with 1ug/mL of RNase A (EN0531) in low salt for 30 min at 37 C after fixation and permeablization but before antibody staining.

### Immunocytochemistry

After washing 1x with DPBS, cultures were fixed for 7 min with 4% paraformaldehyde before blocking for 1 hr in DPBS 0.2% Triton X-100, 5% goat serum. Antibodies are incubated in blocking buffer overnight at 4C (MX1, Cell Signaling, 37849, 1:500). Cultures are washed 3x with DBPS before adding secondary antibodies 1:1000 and Hoechst 1:1000 diluted in DPBS. After 30 min, cultures are washed 3x with DPBS and stored in 0.5% NaN^3^.Images were collected on the Molecular Devices Image Express Confocal HT.ai or IN Cell Analyzer 6000. Images were analyzed using CellProfiler.

### Lentivirus generation

Lentivirus was generated as described^36^. Briefly, HEK-293T were plated to achieve 80-95% confluence after 24 hrs. For 2 mL of media, 1 ug each of transfer plasmid and third generation packaging mix were mixed with 100 uL Optimem (Gibco, 31985088) and 12 uL TransIT-Lenti Transfection Reagent (Mirusbio, MIR 6604). Transfection mix was incubated for 10 min at room temperature before adding to HEK-293T cells. After 48 hrs, medis was collected and filtered through a 0.45 um PVDF filter. Lentivirus Precipitation solution (Alstem, VC125) was used per manufacturer’s protocols.

iPSCs were infected with virus at time of passage and allowed 48 hrs for expression before selection with 1 ug/mL puromycin for two passages or until >95% BFP positive populations were achieved. For CRISPR screening libraries, cells were transduced at an MOI of 0.3 to avoid double sgRNA transduction within the same cell.

### Genome-wide CRISPRi screen

For performing the genome-wide screen, seven sublibraries (CRISPRi_v2) were cloned into the pMK1334 background (Addgene, 127965) as described previously^36,37^. For each sublibrary, iPSCs were transduced at an MOI of 0.3 with 1000x guide coverage. For all maintenance of iPSCs with CRISPRi libraries including routine passaging, freezing, and differentiation, cells were maintained at at least 1000x coverage.

For IFIT1 screening, each sublibrary of iPSCs was plated to eight 15 cm plates at 10M iPSCs per plate and differentiated as described above with TMP to induce CRISPRi activity starting at d0. On day 8 of differentiation, iPS-Microglia were treated with 10 pM IFNβ for 24 hrs. On day 9, iPS-Microglia were lifted with TrypLE for 10 min at 37C and quenched with TBS + 3% BSA + 0.5 uM EDTA. After lifting, cells were stained with Zombie Red Fixable Viability kit (Biolegend, 423110, 1:200) for 10 min at 4C. Because formaldehyde-based fixation interrupts subsequent genomic DNA isolation and PCR, iPS-Microglia were fixed with zinc fixation buffer (0.1M Tris-HCl, pH 6.5, 0.5% ZnCl^2^, 0.5% Zn Acetate, 0.05% CaCl^2^) overnight at 4C. Following fixation, cells were stained with IFIT1 in TBS + 3% BSA + 0.5 uM EDTA for 30 min at 4C. Antibody was washed off in TBS + 3% BSA + 0.5 uM EDTA and cells were strained through a 20 um filter to remove clumps. For each library ≥80M iPS-Microglia were sorted using BD FACSAria Fusion flow cytometer to ensure >1000x coverage of guides in both high and low IFIT1+ populations. Cells were gated on live, single cells, and sorted into top 30% and bottom 30% of IFIT1 expression. After sorting, genomic DNA was isolated with NucleoSpin Blood L kit (Macherey-Nagel, 740954.20). sgRNA casettes were isolated and amplified as previously described^36^ and sequenced on Illumina HiSeq-4000.

### CRISPRi screen analysis

CRISPRi screening libraries were analyzed as previously described^38^. Raw sequencing results were mapped with ‘sgcount’ (https://github.com/noamteyssier/sgcount) and ‘crispr_screen’ (https://github.com/noamteyssier/crispr_screen) was used to perform differential expression analysis using a ‘drop first’ weighting strategy and Benjamini-Hochberg multiple hypothesis correction. Significant guide enrichment was calculated for each sublibrary independently based on the non-targeting guides within that library. For display of all seven sublibraries within the same plot, log2FC enrichment of guide frequencies in high vs low samples were normalized to the standard deviation of non-targeting guides for that sublibrary.

To parse known screen hits, Google Gemini 2.5 pro was used to generate a prompt for Gemini 2.5 deep research. Full prompt is provided in Supplemental Table 4. In brief, Gemini was prompted to determine the strength of information linking each gene hit to interferon signaling by grouping into one of three categories.

Category 1: Canonical interferon signaling genes that directly impact the signal transduction downstream of type I interferon.Typically includes genes well-documented in the canonical JAK-STAT pathway, such as JAK kinases, STAT transcription factors, IRFs, and other well-recognized type I interferon signaling components (e.g., TYK2, ISGF3, etc.).

Category 2: Genes with any paper or limited evidence suggesting they may impact interferon signaling, viral response, or nucleic acid sensing. This includes genes where there is some experimental or in silico evidence linking them to interferon signaling or antiviral pathways, but where the evidence is either indirect, limited in scope, or not as widely confirmed.

Category 3: Genes with no published impact on interferon signaling. No significant literature, gene ontology annotation, or experimental evidence that directly ties them to type I interferon signaling or antiviral mechanisms.

### Reverse-transcriptase quantitative polymerase chain reaction

For independent validation of screening results, individual sgRNAs were added to iPSCs as described above in lentivirus production. For samples with in-well non-targeting control, BFP expression was swapped for mApple expression as previously described^7^. sgRNA sequences available in Supplemental Table 6.

To assess level of RNA knockdown for independent lines, RNA was extracted using the Quick-RNA Microprep Kit (Zymo, R1050). cDNA was generated using SensiFAST cDNA Synthesis Kit (Meridian Bioscience, BIO-65053). For quantitative PCR, SensiFAST SYBR Lo-ROX 2X Master Mix (Bioline; BIO-94005) was used with custom qPCR primers (Supplemental Table 6) from Integrated DNA Technologies. Amplification was quantified using the Applied Biosystems QuantStudio 6 Pro Real-Time PCR System using QuantStudio Real Time PCR software (v.1.3). Expression fold changes were calculated using the ΔΔCt method, normalizing to housekeeping gene *GAPDH*.

### RNA-sequencing

iPS-Microglia were treated with 100 pM IFNβ on day 8 of differentiation. On day 9, iPSCs were lifted with TrypLE for 10 min at 37C. RNA was isolated with Quick-RNA Miniprep Kit (Zymo, R1055). For determination of IFNβ responsive genes, QuantSeq 3’ mRNA-Seq Library Prep Kit for Illumina FWD (Lexogen, 015UG009V0252) was used to prepare RNA-sequencing libraries per the manufacturer’s instructions. Library amplification was performed with 11 PCR cycles. For analysis of differentiation expression after CNOT10 knockdown, PIP-seq 3’ Single-Cell RNA kit v3.0 was used per manufacturer’s instructions. Libraries were sequenced on Illumina NextSeq-2000. Transcripts were aligned to the human reference genome (GRCh38, release 42) using Salmon v.1.4.Transcripts were mapped to genes and quantified using tximeta.

Cells were filtered to only include those with a minimum count of 2,500. 5,000 of the most highly variable genes were selected with a minimum shared count of 10. Cells were then normalized to a target sum of 10,000 and log transformed. The highly variable genes were used to perform principal component analysis. Next, nearest neighbors were calculated using the top 15 principal components, selecting the 15 nearest neighbors. All single-cell filtering, transformation, and dimensionality reduction was conducted using ‘scanpy’^39^.

To measure perturbation effect in cells, we first calculated a perturbation signature for each cell by searching for the 20 nearest neighbors from non-targeting cells (NTC), and then removing technical variation so that the perturbation-specific effect can be revealed. This was done using the CalcPerturbSig() function from the ‘Mixscale’ developed by the Satija lab^40^ and results were stored in the PRTB assay in ‘Seurat’^41,42^. We then conducted differential expression analysis between CNOT10 knockdown (KD) and NTC cells using a Wilcoxon rank-sum test, using the ‘presto’ package^43^ (https://github.com/immunogenomics/presto) as used in Seurat v5 but implemented via a custom version of the ‘Mixscale’ package (https://github.com/reetm09/mixscale). Our implementation modified the original approach in identifying differentially expressed genes (DEGs) by applying Benjamini-Hochberg correction for multiple hypothesis testing instead of Bonferroni correction. DEGs with a p-value < 0.05 after adjustment were selected as significant DEGs.

We used the top 500 significant DEGs to quantify the magnitude of expression shift of each cell from NT cells due to perturbation. This was done by building perturbation vectors for an equal number of cells in KD and control, where the length of each vector was equal to the number of identified DE genes, with expression values from the PRTB assay. By calculating the mean difference in perturbation vectors between KD and NT cells, we projected each cells’ perturbation vector onto the difference vector to obtain perturbation scores. Scores were standardized by calculating the mean and standard deviation from NT cells. These standardized Mixscale scores were then used to weight cells in a negative binomial regression DE analysis, using the Run_wmvRegDE() function, such that cells with a stronger perturbation response had a greater contribution in the identification of DEGs.

This final list of DEGs was then used for downstream analyses including volcano plots and functional interpretation.

## Author Contributions

AM and MK designed and conducted and analyzed the overall research, created figures, and wrote the manuscript. AM and VCC performed the CRISPRi genome-wide screen. AM, VCC, NT, and WL analyzed the CRISPRi genome-wide screen. OT, KL, VCC, VH, and TT generated independent knockdown lines. AM and RM analyzed RNA-sequencing data. All authors reviewed and approved the final manuscript.

## Competing Interests

MK is a co-scientific founder of Montara Therapeutics and serves on the Scientific Advisory Boards of Montara Therapeutics, Engine Biosciences, Alector and Neurocrine, and is an advisor to Modulo Bio and Theseus Therapies. MK is an inventor on US Patent 11,254,933 related to CRISPRi and CRISPRa screening, and on a US Patent application on in vivo screening methods.

## Acknowledgements

We acknowledge all members of the Kampmann lab for their technical expertise and advice. We thank Caroline Mrejen and the Innovation Core at the Weill Institute for Neurosciences for sharing their microscopy expertise and Sarah Elmes and the Laboratory for Cell Analysis for sharing flow cytometry and FACS expertise. This work was supported by Alzheimer’s Association grants AARF-22-973222 (AM), ZEN-22-969903 (MK), and ADSF-21-831212-C (MK), Larry L. Hillblom Foundation grant 2022-A-016-FEL (AM), National Science Foundation Graduate Research Fellowship under Grant No. 2034836 (OT), and Chan Zuckerberg Initiative grant CP2-1-0000000332 (MK).

## Figures and Figure Legends

**Supplemental Figure 1.**
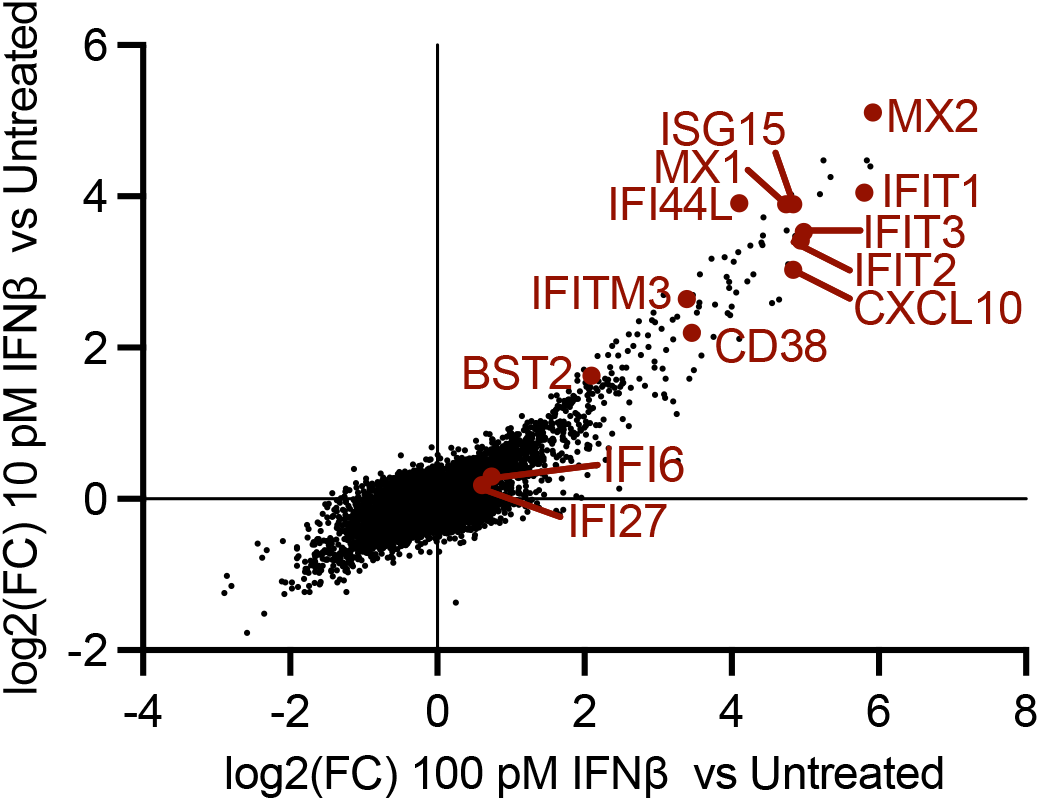
Comparison of differential expression in 10 pM vs 100 pM IFNb treatment. Differentially expressed genes in 100 pM IFNβ treated iPS-Microglia versus untreated are displayed on X axis. Differentially expressed genes in 10 pM IFNβ treated iPS-Microglia versus untreated are displayed on Y axis. Conserved ISGs are denoted in dark red.

**Supplemental Figure 2.**
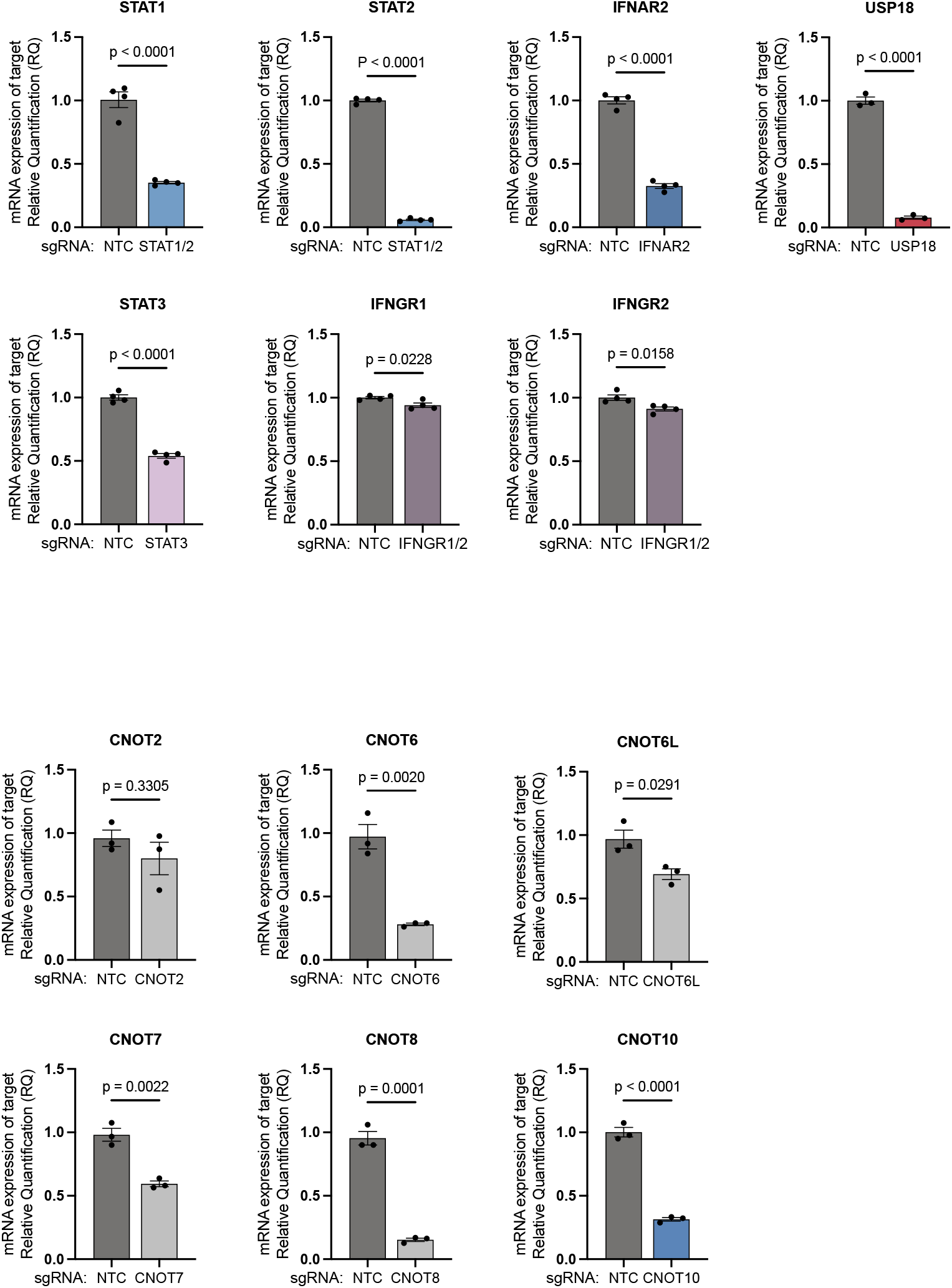
mRNA knockdown in individual knockdown lines. Quantitative reverse transcriptase polymerase chain reaction (RT-qPCR) was performed for each individual knockdown line compared to a non-targeting control (NTC) sgRNA. n=3-4 biological replicates. Bars show mean normalized to the NTC +/-standard error, T-test.

## REFERENCES

1. Baruch, K. et al. Aging. Aging-induced type I interferon signaling at the choroid plexus negatively affects brain function. Science 346, 89–93 (2014).

2. Rexach, J. E. et al. Tau Pathology Drives Dementia Risk-Associated Gene Networks toward Chronic Inflammatory States and Immunosuppression. Cell Rep 33, 108398 (2020).

3. Magusali, N. et al. A genetic link between risk for Alzheimer’s disease and severe COVID-19 outcomes via the OAS1 gene. Brain awab337 (2021) doi:10.1093/brain/awab337.

4. Roy, E. R. et al. Type I interferon response drives neuroinflammation and synapse loss in Alzheimer disease. J Clin Invest 130, 1912–1930 (2020).

5. Escoubas, C. C. et al. Type-I-interferon-responsive microglia shape cortical development and behavior. Cell (2024) doi:10.1016/j.cell.2024.02.020.

6. Zimmerman, A. W. et al. Cerebrospinal fluid and serum markers of inflammation in autism. Pediatr Neurol 33, 195–201 (2005).

7. Teter, O. M. et al. CRISPRi-based screen of autism spectrum disorder risk genes in microglia uncovers roles of ADNP in microglia endocytosis and synaptic pruning. Mol Psychiatry 1–18 (2025) doi:10.1038/s41380-025-02997-z.

8. Da Mesquita, S. et al. Aging-associated deficit in CCR7 is linked to worsened glymphatic function, cognition, neuroinflammation, and β-amyloid pathology. Sci Adv 7, (2021).

9. Deczkowska, A. et al. Mef2C restrains microglial inflammatory response and is lost in brain ageing in an IFN-I-dependent manner. Nat Commun 8, 717 (2017).

10. Buckley, M. T. et al. Cell-type-specific aging clocks to quantify aging and rejuvenation in neurogenic regions of the brain. Nat Aging 3, 121–137 (2023).

11. Wang, Q. et al. Brain derived β-interferon is a potential player in Alzheimer’s disease pathogenesis and cognitive impairment. Alzheimer’s Research & Therapy 16, 271 (2024).

12. Deczkowska, A., Baruch, K. & Schwartz, M. Type I/II Interferon Balance in the Regulation of Brain Physiology and Pathology. Trends in Immunology 37, 181–192 (2016).

13. Roy, E. R. et al. Concerted type I interferon signaling in microglia and neural cells promotes memory impairment associated with amyloid β plaques. Immunity 0, (2022).

14. Roy, E. R., Li, S., Saroukhani, S., Wang, Y. & Cao, W. Fate-mapping and functional dissection reveal perilous influence of type I interferon signaling in mouse brain aging. Mol Neurodegener 19, 48 (2024).

15. Udeochu, J. C. et al. Tau activation of microglial cGAS-IFN reduces MEF2C-mediated cognitive resilience. Nat Neurosci 26, 737–750 (2023).

16. Xie, X. et al. Activation of innate immune cGAS-STING pathway contributes to Alzheimer’s pathogenesis in 5×FAD mice. Nat Aging 3, 202–212 (2023).

17. Sliter, D. A. et al. Parkin and PINK1 mitigate STING-induced inflammation. Nature 561, 258–262 (2018).

18. Interferon beta-1b is effective in relapsing-remitting multiple sclerosis. I. Clinical results of a multicenter, randomized, double-blind, placebo-controlled trial. The IFNB Multiple Sclerosis Study Group. Neurology 43, 655–661 (1993).

19. Jacobs, L., O’Malley, J., Freeman, A. & Ekes, R. Intrathecal interferon reduces exacerbations of multiple sclerosis. Science 214, 1026–1028 (1981).

20. Ejlerskov, P. et al. Lack of Neuronal IFN-β-IFNAR Causes Lewy Body- and Parkinson’s Disease-like Dementia. Cell 163, 324–339 (2015).

21. Yang, X. et al. Functional characterization of Alzheimer’s disease genetic variants in microglia. Nat Genet (2023) doi:10.1038/s41588-023-01506-8.

22. Hasselmann, J. et al. Development of a Chimeric Model to Study and Manipulate Human Microglia In Vivo. Neuron (2019) doi:10.1016/j.neuron.2019.07.002.

23. Mancuso, R. et al. Stem-cell-derived human microglia transplanted in mouse brain to study human disease. Nat. Neurosci. 22, 2111–2116 (2019).

24. Olah, M. et al. Single cell RNA sequencing of human microglia uncovers a subset associated with Alzheimer’s disease. Nature Communications 11, 6129 (2020).

25. Dräger, N. M. et al. A CRISPRi/a platform in human iPSC-derived microglia uncovers regulators of disease states. Nat Neurosci (2022) doi:10.1038/s41593-022-01131-4.

26. Liu, S.-Y., Sanchez, D. J., Aliyari, R., Lu, S. & Cheng, G. Systematic identification of type I and type II interferon-induced antiviral factors. Proceedings of the National Academy of Sciences 109, 4239–4244 (2012).

27. Yamada, T., Horisberger, M. A., Kawaguchi, N., Moroo, I. & Toyoda, T. Immunohistochemistry using antibodies to alpha-interferon and its induced protein, MxA, in Alzheimer’s and Parkinson’s disease brain tissues. Neurosci Lett 181, 61–64 (1994).

28. Kazakov, A. S. et al. Interferon-β Activity Is Affected by S100B Protein. Int J Mol Sci 23, 1997 (2022).

29. Du, H. et al. RNF219 interacts with CCR4–NOT in regulating stem cell differentiation. Journal of Molecular Cell Biology 12, 894–905 (2020).

30. Raisch, T. et al. Reconstitution of recombinant human CCR4-NOT reveals molecular insights into regulated deadenylation. Nat Commun 10, 3173 (2019).

31. Zhou, J. et al. LilrB3 is a putative cell surface receptor of APOE4. Cell Res 33, 116–130 (2023).

32. Mauxion, F., Prève, B. & Séraphin, B. C2ORF29/CNOT11 and CNOT10 form a new module of the CCR4-NOT complex. RNA Biol 10, 267–276 (2013).

33. Höpfler, M. et al. Mechanism of ribosome-associated mRNA degradation during tubulin autoregulation. Molecular Cell 83, 2290-2302.e13 (2023).

34. Heraud-Farlow, J. E. et al. GGNBP2 regulates MDA5 sensing triggered by self double-stranded RNA following loss of ADAR1 editing. Sci Immunol 9, eadk0412 (2024).

35. Gordon, D. E. et al. A Quantitative Genetic Interaction Map of HIV Infection. Mol Cell 78, 197-209.e7 (2020).

36. Tian, R. et al. CRISPR Interference-Based Platform for Multimodal Genetic Screens in Human iPSC-Derived Neurons. Neuron 104, 239-255.e12 (2019).

37. Horlbeck, M. A. et al. Compact and highly active next-generation libraries for CRISPR-mediated gene repression and activation. eLife 5, e19760 (2016).

38. Ramani, B. et al. CRISPR screening by AAV episome-sequencing (CrAAVe-seq) is a highly scalable cell type-specific in vivo screening platform. bioRxiv 2023.06.13.544831 (2024) doi:10.1101/2023.06.13.544831.

39. Wolf, F. A., Angerer, P. & Theis, F. J. SCANPY: large-scale single-cell gene expression data analysis. Genome Biology 19, 15 (2018).

40. Jiang, L. et al. Systematic reconstruction of molecular pathway signatures using scalable single-cell perturbation screens. Nat Cell Biol 27, 505–517 (2025).

41. Hao, Y. et al. Dictionary learning for integrative, multimodal and scalable single-cell analysis. Nat Biotechnol 42, 293–304 (2024).

42. Stuart, T. et al. Comprehensive Integration of Single-Cell Data. Cell 177, 1888-1902.e21 (2019).

43. Korsunsky, I., Nathan, A., Millard, N. & Raychaudhuri, S. Presto scales Wilcoxon and auROC analyses to millions of observations. 653253 Preprint at 10.1101/653253 (2019).

