## Supplemental Table 4 for "Regulators of Interferon-Responsive Microglia Uncovered by Genome-wide CRISPRi Screening"

**Task:** You will receive a list of genes below. For each gene in this list, determine how strongly it is associated with type I interferon signaling (especially canonical interferon signal transduction) and/or antiviral response, and then classify it into one of the following three categories:          
Category 1: Canonical interferon signaling genes that directly impact the signal transduction downstream of type I interferon. Typically includes genes well-documented in the canonical JAK-STAT pathway, such as JAK kinases, STAT transcription factors, IRFs, and other well-recognized type I interferon signaling components (e.g., TYK2, ISGF3, etc.).
Category 2: Genes with any paper or limited evidence suggesting they may impact interferon signaling, viral response, or nucleic acid sensing. This includes genes where there is some experimental or in silico evidence linking them to interferon signaling or antiviral pathways, but where the evidence is either indirect, limited in scope, or not as widely confirmed.          
Category 3: Genes with no published impact on interferon signaling. No significant literature, gene ontology annotation, or experimental evidence that directly ties them to type I interferon signaling or antiviral mechanisms.

**Instructions for Classification:**          
Literature Review: Search for each gene’s name, aliases, and standard Gene Symbols in major literature databases (e.g., PubMed, Google Scholar, or references contained in curated resources).          
Check if it is mentioned in the context of type I interferon, JAK-STAT signaling, IRFs, ISGs, antiviral immunity, or viral response research articles.          
Take note of the quality and quantity of evidence: Is it cited in well-known reviews or studies explicitly as part of the core IFN pathway?          
Is there only a single or a small handful of studies linking it tangentially to interferon or antiviral response?          

Database & Ontology Queries: Use gene ontology (GO) resources, such as the Gene Ontology website, and official pathway repositories (e.g., KEGG, Reactome, or WikiPathways) to check for terms like: “type I interferon signaling pathway,” “response to virus,” “immune response,” “JAK-STAT cascade,” etc. If the gene is annotated with GO terms clearly indicating involvement in “type I interferon signaling pathway,” “viral defense,” or “immune response,” prioritize that information.          

Categorization Based on Evidence:         
Category 1 (Canonical IFN signaling): The gene is frequently cited and recognized as a core component of type I IFN signaling (e.g., direct effectors in the JAK-STAT cascade, known IRFs, or well-characterized interferon-stimulated genes).          

Category 2 (Few studies or indirect involvement): The gene appears in at least a few publications linking it to interferon pathways or antiviral response, but it is not widely recognized or directly annotated as a canonical signaling component.

May be implicated in modulating IFN pathways or viral replication in cell-based or organismal studies, but evidence is not robust or is limited to specialized contexts.

Category 3 (No known impact):The gene has no significant mention in the context of interferon biology or viral response in literature or gene ontology/pathway databases.          

Report Format- Return a table or structured data containing:          
Gene Symbol: The official Gene Symbol (e.g., as designated by HGNC).          
Common Aliases: Any synonyms or alternative names.          
Category (1, 2, or 3).          
Evidence Summary: (Optional but recommended) A short note with either a citation or a brief description of the supporting rationale (for categories 1 and 2), or a statement that no evidence was found (for category 3).          

Important Notes:

1. If conflicting data emerges, prioritize recent and well-reviewed publications or authoritative databases.
2. If a gene is widely known to be essential in another closely related pathway (e.g., NF-κB activation) but only loosely tied to IFN signaling, consider whether it truly belongs in Category 2 or is best assigned Category 3.
3. Remember: The objective is to comprehensively classify each of these genes into exactly one category based on the current state of scientific evidence.
